# A genetically encoded adhesin toolbox for programming multicellular morphologies and patterns

**DOI:** 10.1101/240721

**Authors:** David S. Glass, Ingmar H. Riedel-Kruse

## Abstract

Synthetic multicellular systems hold promise for understanding natural development of biofilms and higher organisms^1,2^, as well as for engineering complex multi-component metabolic pathways^2,3^ and materials^4^. However, such efforts will require tools to adhere cells into defined morphologies and patterns, and these tools are currently lacking^1,5,6^. Here we report the first 100% genetically encoded synthetic platform for modular cell-cell adhesion in *Escherichia coli*, which provides control over multicellular self-assembly. Adhesive selectivity is provided by a library of outer membrane-displayed nanobody^7,8^ and antigen peptides with orthogonal intra-library specificities, while affinity is controlled by intrinsic adhesin affinity, competitive inhibition, and inducible expression. We demonstrate the resulting capabilities for rational design of well-defined morphologies and patterns through homophilic and heterophilic interactions, lattice-like self-assembly, phase separation, differential adhesion, and sequential layering. This adhesion toolbox, compatible with synthetic biology standards^9^, will enable construction of high-level multicellular designs and shed light on the evolutionary transition to multicellularity^6,10^.

---

Multicellular organisms display a variety of morphologies (3-dimensional structures) and patterns (spatial distributions of cell types) over multiple length scales using multiple cell types. There is growing interest in synthetic biology^1,2,11–15^ to engineer such multicellular arrangements in order to harness their unique abilities, such as to separate intermediates in increasingly complex metabolic pathways^2,3^ or to program structured living materials^4^ and tissues^16–18^. For example, synthetic circuits were engineered that direct cells on 2D substrates into self-organized ring patterns^12,19^. Such patterns were enabled by tools for cell-cell signaling^12,13, 20, 21^ and differentiation^22, 23^. For natural multicellular organisms the key third tool for directing spatial organization is cell-cell adhesion^6,10^, but comparable synthetic tools are lacking. Synthetic cell-cell adhesion tools have been developed to adhere various cell types^18,24–28^, but have limited use for multicellular engineering due to (1) having limited control over specificity^18,24^, (2) only mediating adhesion among very different cell types (bacterial-to-mammalian)^25^, or (3) having a non-genetic basis requiring chemical modifications that are diluted by growth^26–28^. We propose that a synthetic cell-cell adhesion toolbox should have the following properties: it should be (1) genetically encoded, (2) decoupled from native signaling and adhesion, (3) easily extendable to an arbitrary library of adhesins, (4) tunable in binding strength and binding specificity, and (5) compatible with cell growth and division.

Here we present a synthetic cell-cell adhesion toolbox in *E. coli* that meets these criteria and enables controlled multicellular self-assembly (Fig. 1). We designed this toolbox from three elements: a transcriptional regulator, an outer membrane anchor, and an adhesin library. The regulator for adhesin display is a standard TetR or AraC repressor, controlled by addition of small molecule inducers anhydrotetracycline (ATc) or arabinose (Ara), respectively (Fig. 1a). Outer membrane anchoring is achieved via the intimin N-terminus of enterohemorrhagic E. *coli* (EHEC O157:H7), a reverse autotransporter and surface display system that includes a short N-terminal signal peptide to direct its trafficking to the periplasm, a LysM domain for peptidoglycan binding, and a *β*-barrel for transmembrane insertion^8,24,25^ (Figs. 1a,b). As adhesins we use nanobodies and their antigens^7,8^ (Figs. 1a,b). Nanobodies, the variable domains of camelid heavy-chain antibodies, can be expressed on bacterial surfaces due to their small size (~ 125 amino acids) and stability under a variety of conditions^7^. Their single-domain architecture, in combination with the intimin autotransporter, allows the entirety of a highly specific cell-surface adhesin in a single fusion protein. We hypothesized that two *E. coli* strains displaying a nanobody and its antigen, respectively, would specifically adhere to each other (Figs. 1b,c), that this would allow controlled morphology and patterning of multicellular assemblies (Fig.1c), and that a library of such adhesin pairs could be established (Fig. 1d).

**Figure 1.**
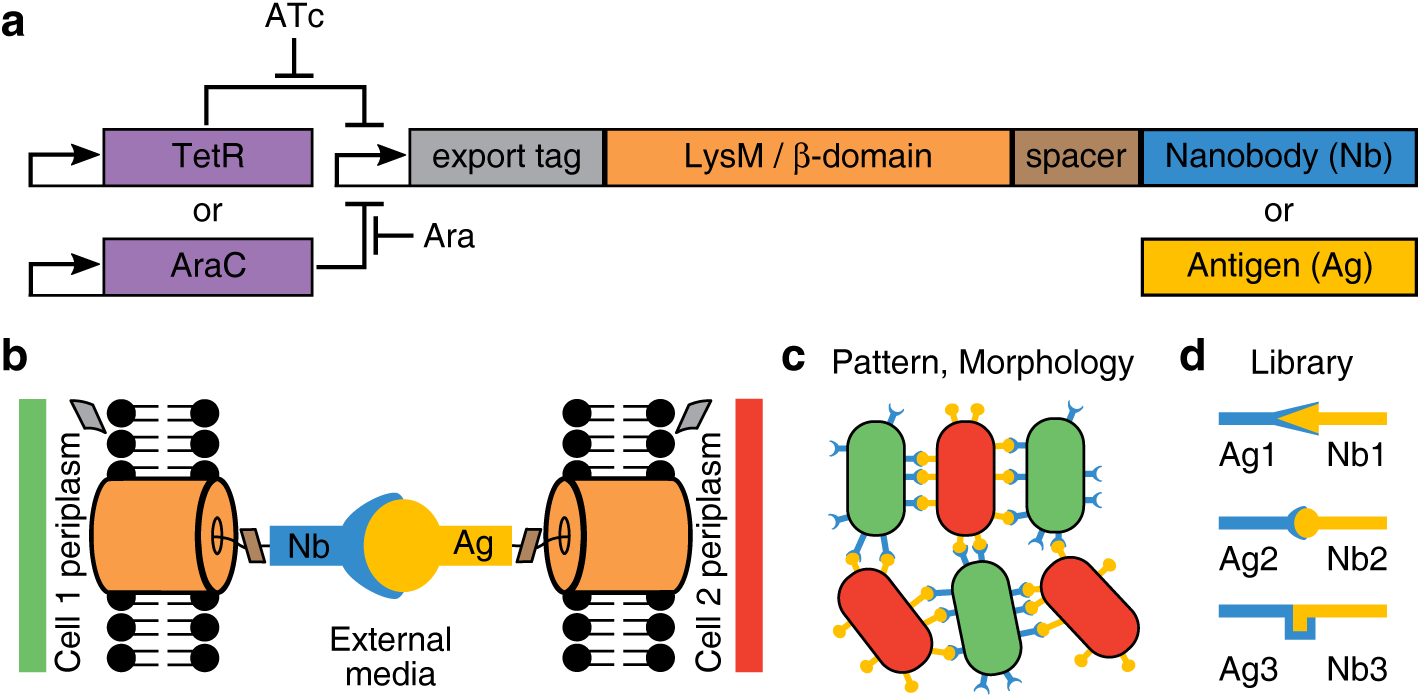
Design of a synthetic cell-cell adhesion toolbox that allows aggregation of multicellular patterned morphologies. **a**, Adhesin constructs consist of a single coding sequence, with a nanobody (Nb) or antigen (Ag) fused to the autotransporter initimin N-terminus from enterohem-orrhagic *E. coli*. A repressor (TetR or AraC), coupled to an inducer (ATc or Ara) that relieves the repression, can be used to control expression levels. **b**, At the periplasm, intimin folds into the outer membrane, displaying the spacer and adhesin (Nb or Ag) outside the cell. **c**, Nb-Ag interactions between cells should mediate adhesion and microscopic, patterned morphologies (here alternating positioning of red and green cells into 3D aggregates). **d**, A library of adhesin pairs can be used to expand adhesion capabilities.

To establish the corresponding adhesin library, we synthesized 51 nanobodies and 8 short antigens (≲ 125 amino acids to match the nanobodies) whose sequences we obtained from the VIB Nanobody Core (see methods). We transformed this intimin fusion library into MG1655 wild type K-12 *E. coli* with a single adhesin per strain. Stationary-phase cultures were left unshaken alone or in mixture with their corresponding Ag- or Nb-expressing partner strain, with cell-cell adhesion detectable by macroscopic aggregation and settling (Figs. 2a,b) within ~ 1 hour (Fig. S1). We quantified this aggregation by measuring optical density (OD_600_) of cells remaining unaggregated in the upper half of the cultures after 24 hours (so as to detect adhesion interactions stringently). We identified three pairs of strains that aggregated and fell out of solution, comprising 6 strains termed Ag1-3 and Nb1-3 for the antigens and corresponding nanobodies, respectively (Fig. 2c). Multiple nanobodies were found for Ag3 (see below). Importantly, no such aggregation occurred in monocultures, in co-cultures without induction by ATc, or in a Null control containing the autotransporter but no adhesin (Fig. 2c).

**Figure 2.**
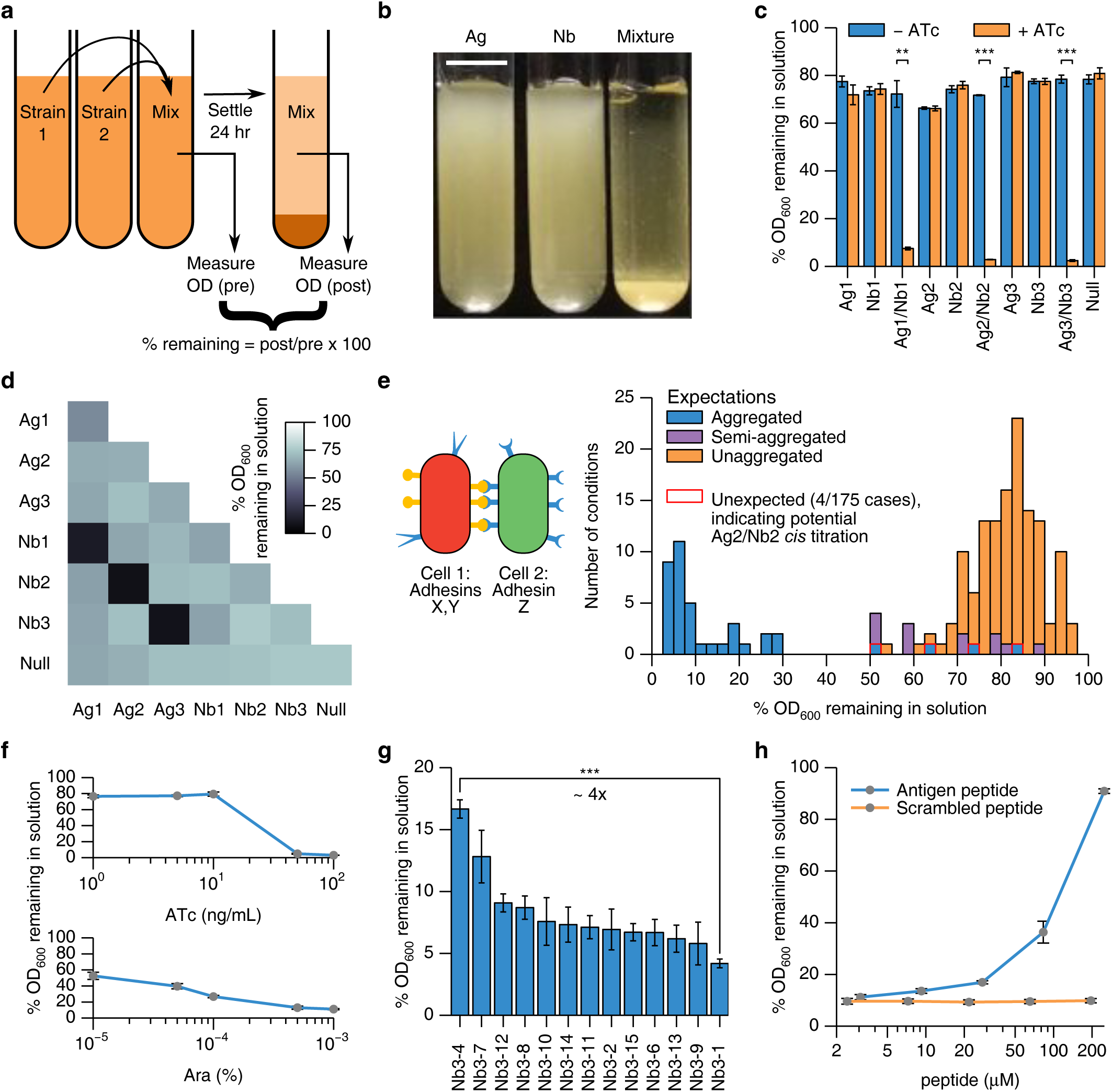
A library of adhesins enables multiple ways of tuning adhesion specificity and strength. **a**, An aggregation assay using optical density (OD_600_) measurements allows the quantification of binding strength and specificity between cells. **b**, Binding in a 1:1 ratio of Ag2:Nb2 cell types leads to macroscopic aggregation and settling. Scale bar: 1 cm. **c**, Aggregating mixtures for three Ag/Nb pairs show significant settling compared to unmixed, uninduced, and no-adhesin (Null) conditions. **d**, Nb/Ag-based adhesion interactions are orthogonal, as strains only aggregate significantly with their designed partner strain (*p* < 0.001 for 2-tailed ¿-test compared to 50% decrease). **e**, Multiple adhesins can be used simultaneously (composability). Twenty-five strains containing all permutations of 5 adhesins (Ag2, Ag3, Nb2, Nb3, Null) on medium-copy (X) and low-copy (Y) plasmids were mixed with the original single-adhesin, medium copy strains (Z). Aggregating cultures (%OD_600_ ≲ 40) behave as expected except for strains containing both Ag2 and Nb2 in the same cell, indicating *cis* interactions for this pair. Semi-aggregated refers to expected mixtures of non-aggregating and (homophilically) aggregating cells.f, Aggregation of Ag2/Nb2 under different concentrations of ATc (top) and Ara (bottom) show induction control. **g**, Aggregation of 13 different Nb3 variant strains (ranked by %OD result) mixed with the Ag3 strain shows adhesion can be controlled by controlling individual Nb/Ag affinity, **h**, Aggregation of Ag2/Nb2 is diminished upon addition of soluble Ag2 peptide (sequence: EPEA) but not of a control peptide (sequence: PEAE). Displayed values are averages for *N* = 3 samples. Error bars: ±1 S.D., *p* < 0.01 (**) or p < 0.001 (***) according to a 2-tailed paired *t*-test.

We tested that this nanobody-antigen library is both orthogonal (adhesins only interact with designed partners) and composable (multiple adhesins function simultaneously within one cell). To demonstrate orthogonality, we assayed aggregation in all pairwise mixtures of the 7 strains used in Fig. 2c (Ag1-3, Nb1-3, Null), and indeed significant reduction in supernatant density only occurred for mixtures of the designed Ag-Nb pairs. To demonstrate composability, we focused on Ag2, Ag3, Nb2, Nb3, and Null strains. We produced a set of 25 strains, each expressing 2 of these 5 adhesins simultaneously, with one adhesin on a low-copy plasmid and the other on a medium-copy plasmid for convenience. We assayed aggregation of each of the 25 strains when mixed with the original 5 strains expressing just one of the 5 adhesins, for a total of 25×5=125 conditions (Fig. 2e left). We also assayed aggregation in unmixed samples of the 25 strains with and without induction by ATc, for an additional 25×2=50 conditions. Of the 125+50=175 total conditions, all but four (97.7%) behaved as expected (Fig. 2e right), aggregating if and only if a nanobody and its corresponding antigen were both present in the mixture (see Figs. S2-S3 for details). The remaining four represent unsuccessful homophilic adhesion (i.e., between like cells) of cells producing both Ag2 and Nb2, which we speculate is due to *cis* titration of Nb2 by the much smaller (4 amino-acid) Ag2 peptide when both are expressed in the same cell (Fig. S2).

We next established the ability to control adhesion strength quantitatively between cells through three independent methods. First, we controlled affinity on a per-cell basis by controlling adhesin expression level (Fig. 2f). Varying inducer concentration over 3 orders of magnitude showed rather digital on/off control over aggregation by ATc and a more graded response but substantial leaky expression for Ara. Second, we controlled affinity on a per-molecule basis by individual Nb-Ag affinity. This we demonstrated using multiple nanobodies against Ag3, which showed a range of binding strengths as indicated by the percentage of cells remaining in solution (Fig. 2g). Third, we used a soluble peptide to competitively inhibit cell-cell adhesion. In particular, soluble Ag2 peptide blocked the formation of aggregates between Ag2 and Nb2 strains in a concentration-dependent manner, while a scrambled-sequence control peptide had no effect (Fig. 2h). The potential *cis* titration of Nb2 by Ag2 mentioned above could provide a fourth method similar competitive inhibition by soluable Ag2.

With this synthetic adhesion toolbox in hand, we explored what multicellular morphologies and spatial patterns could be achieved at a microscopic scale with a single adhesin pair expressed in two strains. We labeled Ag2- and Nb2-expressing strains with constitutive, cytoplasmic fluorescent proteins mRuby2 (red) and sfGFP (green), respectively, leading to extended aggregates with mesh-like patterns of alternating red and green cells (Fig. 3a). This red-green binding specificity is statistically significant when quantifying the number of nearest neighbors (Fig. 3b) and the nearest-neighbor matrix, or conditional probability table (Fig. 3c, Supplementary Discussion).

**Figure 3.**
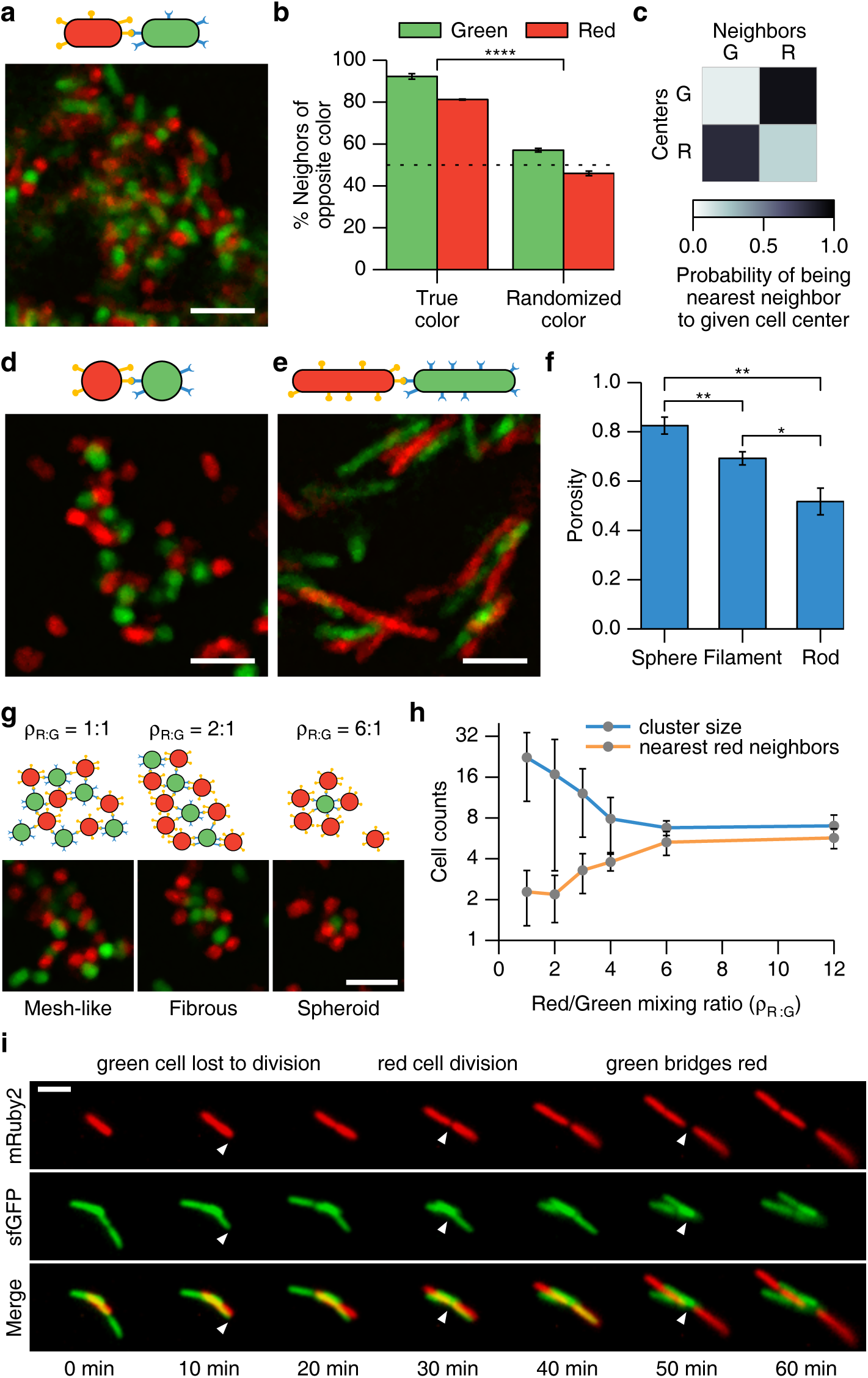
The adhesion toolbox enables self-assembly of multicellular aggregates with defined morphologies and precise lattice-like patterning, and is compatible with cell growth and division. **a**, A single confocal z-slice from an Ag2/Nb2 aggregate. Ag2 cells express cytoplasmic mRuby2 (Red); Nb2 cells express cytoplasmic sfGFP (Green). **b**, Quantification of 3D confocal stacks as in a. Most cells neighboring any given cell are of the opposite cell type. Randomizing cell identities shows approximately uniform number of neighboring cell types (*p* = 1.2 × 10^−6^ for true vs. random coloring by two-tailed *t*-test). Lower percentages for red cell centers is likely due to there being 24 ± 3% more red cells than green cells in these samples (see g). **c**, Conditional probability table for **a**, giving the chance for a cell of a given color (column) being the nearest-neighbor to a cell of another color (row). See Supplemental Discussion. **d**,e, Same as in **a**, but with spherical and filamentous cells, respectively. **f**, Porosity quantification of aggregates from **a**,d,e show significant differences in structure. **g**, Increasing the density ratio *ρ*_R:G_ of spherical cells changes morphology (cluster size and shape) and patterning (nearest red neighbors per green cell) predictably as available binding partners for green cells decreases (schematic, top). **h**, quantification of g. **i**, Time lapse of adhesive co-cultures shows patterning even during cell growth and division (Ag2/sfGFP + Nb2/mRuby2). Scale bars: 5 *μ*m. Displayed values are averages for *N* = 3 confocal stacks. Error bars: ±1 S.D. Asterisks as in Fig. 2 with **** representing *p* < 0.0001.

We then tested whether the aggregate morphology could be modulated with alternative cell morphologies, using a spherical S1 strain (due to *mrdB* mutation^29^, Fig. 3d) or a filamentous strain (MG1655 with membrane stress due to overexpression off of a high-copy plasmid^30^, Fig. 3e). The resulting aggregates differ significantly both in their microscopic porosity (fraction of space not occupied by cells, Fig. 3f) and in their macroscopic pellet size (Fig. S4). That is, rods (wild type) pack more compactly than filaments, and the spherical S1 mutants (for an undetermined reason) do not even form macroscopic aggregates.

We further tested whether we could control aggregate size and morphology in S1 cells by varying the density ratio *ρ*_R:G_ between two adhering cell types (Fig. 3g). We found a transition (Fig. 3g,h) from large, mesh-like structures (*ρ*_R:G_ ≈ 1 : 1) to more elongated fibrous clusters (*ρ*_R:G_ ≈ 2 : 1) and eventually to small spheroids (p_R:G_ ≈ 6:1). These different morphologies and cluster sizes are expected as the more abundant cell type makes maximal use of the available binding sites around the less abundant cell type by surrounding it. Thus overall, the adhesion toolbox enables production of aggregates with lattice-like patterns and modulation of aggregate size, morphology, and material properties such as porosity.

We also established that our toolbox is compatible with patterning throughout cell growth and division. We tracked small aggregates of exponential-phase cells in a microfluidic chamber^31^ for several hours. We observed cells growing and dividing over multiple cell cycles while also adhering to other cells (Fig. 3i). Pairs of red and green cells bound lengthwise gave rise to multiple generations of daughter cells similarly bound lengthwise, leading to small filaments of two to three cell widths. Absent an adjacent cell of the opposite color, daughter cells separated from the aggregate after division (Fig. 3i left). Conversely, the presence of the opposite cell type maintained daughter cells as part of the aggregate by acting as an adhesive bridge (Fig. 3i right).

Next, we sought to rationally design distinct patterns involving more than one adhesion pair in two cell types by implementing three key canonical patterning processes (Fig. 4a-c): differential adhesion^32^, phase separation^32^, and coaggregation bridging^33^. (1) The differential adhesion mechanism^32^ enables patterning via differences in adhesion strength, wherein all cells bind to all other cells, but those with stronger binding localize to the center of an aggregate and those with weaker binding localize to the periphery. We achieved this (Fig. 4a) with a mixture of four strains producing Ag2 or Nb2 in high or low amounts, and where the high- and low-expressing cells were fluorescently labeled red (mRuby2) and green (sfGFP), respectively. (2) Phase separation, by contrast, can occur in the limit of no adhesion interaction between two groups of cells^32^. We achieved this (Fig. 4b) by generating a blue cell (Cerulean) that is homophilically adhesive through simultaneous expression of Ag3 and Nb3, and then co-mixing this cell with the heterophilically adhering Ag2 (green, Venus) and Nb2 (red, mCherry) cells demonstrated in Fig. 3a. These separated into homophilic (blue) and heterophilic (green/red) phases. (3) Certain natural systems such as dental plaque biofilms exhibit a phenomenon known as coaggregation bridging^33^, in which two otherwise non-interacting cell types adhere indirectly through an intermediate capable of binding both. We achieved this (Fig. 4c) by generating a blue (Cerulean) cell type presenting both Ag2 and Ag3, which binds green (Venus) Nb3 and red (mCherry) Nb2 cells.

**Figure 4.**
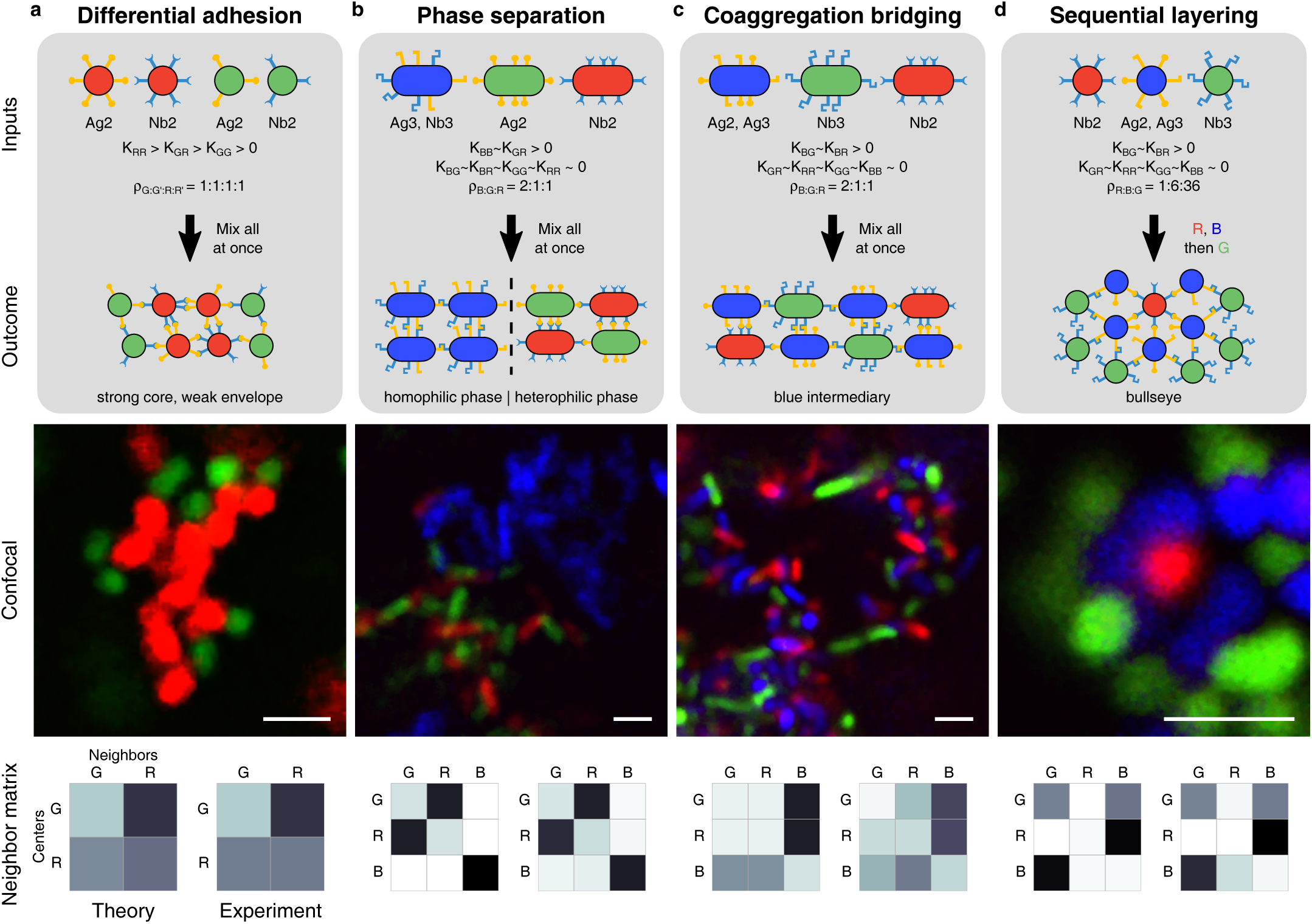
Complex patterns can be rationally designed using the synthetic adhesion toolbox in a combinatorial fashion. **a**, Highly adherent cells (Ag2/mRuby2 + Nb2/mRuby2, 100 ng/mL ATc) and weakly adherent cells (Ag2/sfGFP + Nb2/sfGFP, .0001% Ara) self-assemble through differential adhesion into clusters of red cells surrounded by green cells. **b**, Phase separation between self-adherent homophilic (Ag3/Nb3/Cerulean) and heterophilic (Ag2/Venus + Nb2/mCherry) aggregates. **c**, Coaggregation bridging of non-interacting cell types (Nb3/Venus + Nb2/mCherry) through a doubly adhesive strain (Ag2/Ag3/Cerulean). **d**, Sequential addition of excess binding cells can produce layered “bullseye” clusters. (Nb2/mCherry + Ag2/Ag3/Cerulean, followed by Nb3/Venus). **a-d**, Each panel includes (1) strains and adhesins used, including qualitative estimates of relative association constants (K) and density ratios (*ρ*) between green (G), red (R), and blue (B) cells; (2) mixing protocol; (3) expected patterning outcome and underlying mechanism; (4) typical confocal *z*-slices (Scale bar 2.5 *μ*m); (5) Nearest-neighbor matrices as in Fig. 3c based on a heuristic of pairwise rules using data from Figs. 2,3 (Theory, left); and quantification of confocal images (Experimental, right). These theoretical and experimental neighbor matrices agree quantitatively (see Fig. S5, Supplementary Discussion). Rods are MG1655, spheres are S1. Quantification is averaged over *N* = 3 3D images (**b,c**), 6 clusters of > 15 cells **(a)**, and 9 clusters **(d)**

Finally, we wanted to demonstrate how varying other parameters such as cell shape, density ratio, and timing of culture mixing expands the available pattern space. We achieved this (Fig. 4d) using the same same adhesin set as in Fig. 4c but using spherical cells, mixing the cells sequentially (red, blue, green), and in increasing densities of 1:6:36. This process led to bullseye patterns that differ from Fig. 4c both qualitatively and according to their conditional probability tables (Fig. S5). Note that this bullseye is accomplished solely through cell positioning (via adhesion), in contrast to methods using differentiation and signaling on prepositioned cells^12,19^. Altogether, Fig. 4 demonstrates the adhesion toolbox’s rich patterning capabilities, presenting distinct neighbor-binding matrices and patterning length scales of 2 cells (Fig. 4b heterophilic phase), 3 cells (Fig. 4c,d) and many cells (Fig. 4a,b homophilic phases).

In summary, we established a synthetic cell-cell adhesion toolbox that provides quantified control over key parameters and thereby enables rational programming of versatile multicellular morphologies and patterns through multi-adhesin self-assembly. Toolbox extensions should include sub-cellular localization for symmetry breaking and porting to other cell types including eukaryotes through suitable surface display anchors^34^. This toolbox is compatible with the BioBrick synthetic biology parts assembly standard^9^, enabling higher-level multicellular designs that combine cell-cell signaling^12, 13, 20, 21^, differentiation^22,23^, and logic^11,13^ with adhesin-based control over morphology and patterning. Such implementations should have broad utility for efficient pathway compartmentalization in metabolic consortia engineering^2, 3^ and cell-autonomous morphogenesis in engineered tissues^16–18^ and living materials^4^. Finally, we propose that synthetic, minimal multicellular organisms using these tools should provide bottom-up insights into natural development and the evolutionary transition to multicellularity^6,10^.

## Methods

### Strains and sequences

The two parent strains used in this study, and which were obtained from the Coli Genetic Stock Center (CGSC), are MG1655 (CGSC #6300) and S1 (CGSC #6338). The intimin display system was originally obtained as an IPTG-inducible high copy plasmid termed pNeae2 (and anti-GFP derivative termed pNVgfp) from Luis Ángel Fernández^25^. The Tet-expression and Ara-expression plasmids were con by synthesizing (using IDT’s gBlock service) a restriction site-free version of the intimin N terminus and cloning via BioBrick suffix assembly into iGEM part numbers BBa_K145279 and BBa_I0500, respectively, or into pNeae2 for filamentous strains. Sequences for the nanobody/antigen library were obtained from the VIB Nanobody Core. These were then either synthesized as IDT gBlocks and incorporated into the Tet and Ara expression plasmids via BioBrick suffix assembly or both synthesized and cloned into the Tet expression plasmid by Twist Bioscience. A full list of the nanobody/antigen sequences is available in Supplementary Table S1. Fluorescent protein sequences were obtained from S. DePorter (mRuby2), P. Subsoontorn (sfGFP) or N. Cira (Venus, mCherry, Cerulean) and cloned into various constitutive expression plasmids (pSB1C3, pSB3K3, pSB4A3 with pλ or BBaJ23100, BBa_B0034 expression) by BioBrick assembly (see Supplemental Table S1).

### Aggregation assays

Cultures were grown overnight at 37°C while shaking at 300 rpm in 7 mL LB + 100 ng/mL ATc (if induced and unless noted otherwise) for 24 hours (to ensure stationary phase and consistent final density across samples). Filamentous strains (using pNeae2) were induced using 100 *μ*M isopropyl *β*-D-1-thiogalactopyranoside (IPTG). Cultures were then vortexed briefly and mixed 1:1 with other strains in deep 96-well plates. Samples of 100 *μ*L were taken from the mixtures immediately following mixture and 24 hours later, to ensure equilibrium, from the top ~25% of the well (“supernatant”). Samples were transferred to 96-well assay plates and OD_600_ was measured on a Tecan infinite M1000 plate reader.

### Peptides

EPEA and PEAE peptides were synthesized by Genscript at > 95% purity. The lyophilized peptides were resuspended in water, and their concentration was quantified on a NanoDrop One using the A205/31 method.

### Aggregation time lapses

Cultures were grown and mixed as above, and then transferred to 10 mL clear plastic test tubes, taped to a black felt background with an overhead fluorescent lamp for a dark field effect. Samples were photographed on a Nexus 5X smartphone using the TimeLapse Video Recorder app. Quantification was done in FIJI (Fiji is just ImageJ) by subtracting grayscale values of the upper one third of the test tubes minus neighboring test tubes in the same image containing only media.

### Microscopy

Epifluorescence was performed on a Leica DMI6000B microscope using the GFP and TX2 filter sets, along with brightfield images, and a 40× 0.6 NA objective. Confocal microscopy was performed on a Leica DMRXE microscope using a 63× 1.2 NA water-immersion objective with excitation of 488 nm for sfGFP, 496 nm for Venus, 543 nm for mCherry or mRuby2, and 458 nm for Cerulean. Emission ranges for confocal were manually adjusted to maximize signal and avoid bleed-through. All confocal images were taken after allowing 600 - 1200 *μ*L mixtures to settle in 1.5 mL microcentrifuge tubes for approximately 1 hour. For each sample, approximately 20 of aggregate was extracted from the bottom of the tube using a wide-orifice pipette tip, transferred to a double-sided tape microscope slide chamber, covered with a coverslip, and sealed with Thomas Lubriseal stopcock grease. Sections varying from 20 - 200 *μ*m were imaged by confocal microscopy. Bleed-through from blue to green channels was corrected in 3-color images by subtracting the blue channel from the green channel. For display purposes, a 2-pixel median filter was applied to images in Figs. 3 and 4, and brightness/contrast were adjusted in FIJI for entire images to assure channels appear similar.

### Microscopic time lapse

Overnight cultures were grown for 16 hours, backdiluted 1:1000, grown for 2 hours shaken at 37°C, mixed 1:1 (1.2 mL total), and allowed to settle within a 1.5 mL test tube at 37°C for an additional 2 hours. Using a sterile syringe, ~ 100*μ*L were slowly transferred from the middle of the tube to a microfluidic device made from a layer of polydimethylsiloxane (PDMS) over a glass cover slip, containing an inlet, outlet, and 2 mm × 12 mm × 0.1 mm chamber^31^. The chamber was connected using sterilized steel pins and tubing to two reservoirs of media (3 mL and 2.9 mL) and imaged using epifluorescence in a humidified, 37°C chamber every 5 minutes for 6 hours. Brightness and contrast were automatically adjusted in FIJI for each frame to help identify cells. StackReg ImageJ plugin was used to maintain orientation of cells in Fig. 3i.

### Nearest neighbor quantification

Confocal z-stacks were analyzed using the Bitplane Imaris software package after subtracting the blue channel from the green channel to remove bleed-through on 3-color images, and individual cell centroids were identified using the “surfaces” tool. No filtering or brightness/contrast adjustment was performed in advance. Centroids were analyzed using a custom python script and the scipy NearestNeighbors API. For spherical cells, cell movement between z-stacks precluded automated centroid detection, so centroids were detected in 2D slices either by Imaris as above or by multi-tracker point tool in FIJI, following which the centroids were quantified as above. All code is freely available upon request.

### Porosity quantification

To measure porosity (fraction of volume not occupied by cells), 3D confocal stacks of aggregates were processed in FIJI as follows. First, a 2-pixel median filter was used, and then the images were thresholded using Phansalkar auto local threshold with default parameters. This was done separately for each color channel, and then the color channels were summed. The porosity is reported as one minus the average value of pixels in the thresholded, channel-summed, 3D image stack.

**Supplementary Information** is available in the online version of this paper.

## Acknowledgements

Luis Ángel Fernández provided plasmids pNeae2 and pNVgfp. Gholamreza Hassanzadeh at the VIB Nanobody Core supplied the sequence information for all other nanobodies and antigens. The authors thank X. Jin, A. Spormann, D. Endy, K.C. Huang, H. Kim, N. Cira, A. Keating, and N. Young for helpful discussions, as well as the Spormann and Quake labs for access to their imaging resources. Support was provided by a Stanford Bio-X Bowes fellowship and the American Cancer Society (RSG-14-177-01).

## Author contributions

DSG and IHRK jointly conceived the project and wrote the paper, DSG performed experiments and analysis.

### Author Information

The authors declare no competing financial interests. Correspondence and requests for materials should be addressed to IHRK.

